# A unified peptide array platform for antibody epitope binning, mapping, specificity and predictive off-target binding

**DOI:** 10.1101/2022.06.22.497251

**Authors:** Cody Moore, Anna Lei, Patrick Walsh, Olgica Trenchevska, Gaurav Saini, Theodore M. Tarasow, Mohan Srinivasan, David Smith, Matthew P. Greving

## Abstract

Therapeutic antibody efficacy is largely determined by the target epitope. In addition, off-target binding can result in unanticipated side-effects. Therefore, characterization of the epitope and binding specificity are critical in antibody discovery. Epitope binning provides low-resolution of an antibody epitope and is typically performed as a cross-blocking assay to group antibodies into overlapping or non-overlapping bins. Epitope mapping identifies the epitope with high resolution but requires low throughput methods. In addition to binning and mapping, there is a need for a scalable and predictive approach to reveal off-target binding early in antibody discovery to reduce the risk of in vivo side effects. Peptide microarrays are an information-rich platform for antibody characterization. However, the potential of peptide microarrays in early-stage antibody discovery has not been realized because they are not produced at the scale, quality and format needed for reliable high-throughput antibody characterization. A unified, peptide library platform for high-resolution antibody epitope binning, mapping and predictive off-target binding characterization is described here. This platform uses highly scalable array synthesis and photolithography to synthesize more than 3 million addressable peptides. These arrays conform to a microplate format and each synthesis is qualified with mass spectrometry. Using this platform, a scalable approach to early-stage epitope and specificity characterization, with prediction of off-target interaction(s), is demonstrated using a panel of anti-HER2 monoclonal antibodies. This study highlights the prospect of this platform to improve antibody discovery productivity by generating epitope and specificity information much earlier with potentially hundreds of antibody clones.

## Introduction

Since their advent in the mid 1970s, monoclonal antibodies have become increasingly important molecules for use in research and as therapeutics. Major advancements in the capability to generate, select and sequence diverse antibodies for therapeutic discovery have been realized with B-Cell hybridoma technology, transgenic mice, protein display platforms and next-generation sequencing.^1–8^ Because of these developments, several transformative antibody therapeutics have been discovered and approved by the FDA.^9^ However, in antibody discovery, the corresponding early-stage epitope and specificity characterization platforms used to identify promising leads remain relatively unchanged. Thus the advancements in antibody library generation, selection and sequencing have outpaced the advancements in the ability to comprehensively characterize the binding properties of selected clones early in the process.^10^ This gap between the high number of selected and genetically sequenced antibody clones, and the relatively low number of functionally characterized clones, limits the discovery of novel high-quality antibodies for research and therapeutic applications. In fact, irreproducibility of results and large study failures have been attributed to limited antibody characterization.^10–11^ Recently, major efforts have been initiated to standardize and advance antibody performance metrics.^12^ New platforms that fill the gap between the high number of potentially valuable antibody clones that can be identified, and the limited ability to comprehensively characterize those antibodies, could significantly advance the pace and reliability of antibody-based research and discovery.

Particularly in therapeutic antibody discovery, a platform that can characterize the entire specific and non-specific binding profile in the early stages could improve the productivity of antibody discovery.^13^ Epitope-level antibody characterization is critical because the epitope largely determines antibody activity.^14^ Epitope binning of antibody clones based on their binding to similar or distinct regions on the target is one important measurement because clones that bind distinct epitopes can have very different efficacy and toxicity profiles.^15–25^ Traditional epitope binning assays are performed using label-free biosensors in a competitive assay format that for N number of clones to be binned, at least N^2^/2 pairwise measurements are required.^22, 26–29^ While epitope binning classifies antibody clones based on competition for similar or distinct target binding regions, it does not resolve the identity of the antibody epitopes.^25, 30–32^ Also, pairwise competition assays can produce binning false positives or negatives through several mechanisms including distinct but proximal antibody binding sites and multimeric targets.^30^ Epitope mapping aims to provide position-level resolution of antibody epitopes that are not resolved with epitope binning information. Epitope mapping is another important aspect of therapeutic antibody development because antibodies that have position-level epitope differences can have differences in efficacy, target selectivity, off-target interactions and pan-species reactivity.^17, 21, 30, 33, 34^ Epitope mapping is currently performed using one or more approaches that include X-ray crystallography,^17, 35^ site-directed mutagenesis,^36–38^ shotgun mutagenesis,^39^ mass spectrometry,^35, 40–42^ phage or other display platforms,^16, 33, 43–49^ and solution or array-based peptide libraries.^14, 50–56^ Due to their resource-intensive nature, current high-resolution epitope mapping approaches are limited to a few antibody clones when compared to the total number of potentially valuable clones that can be identified in an antibody screen. Solution or array-based overlapping peptide libraries have the potential for high-throughput epitope mapping but require some prior knowledge about the epitope(s) for their design. Also, typical overlapping peptide libraries have limited sequence coverage that does not include position-level coverage of mutations, deletions, and truncations important to determine antibody tolerance to mutations, predict off-target interactions, and identify pan-species epitopes.^57^ Epitope mapping with *in situ* synthesized peptide arrays can offer comprehensive epitope sequence space coverage with the advantage of an information-rich binding readout for all peptides on the array. Also, peptide arrays are an attractive and unique platform for epitope binning and mapping due to their compatibility with biofluids.^58^ This biofluid compatibility enables direct characterization of immunized animal samples to identify particularly successful immunization strategies from parallel immunizations within an antibody discovery campaign. However, current commercially available *in situ* peptide array production does not scale to the potentially thousands of arrays required for large-scale antibody discovery and are not typically in an automation compatible format.^52, 59, 60^ In addition, current *in situ* synthesized peptide arrays do not offer comprehensive analytical assessment of synthesis yield to ensure the per-synthesis quality required for reliable antibody characterization.

Binding specificity is another important antibody characteristic in the discovery and development of monoclonal antibodies. Off-target interactions arising through CDR-directed binding or any other high-affinity interactions can give rise to undesired outcomes including false positives in research and therapeutic side-effects.^61, 62^ Currently, a scalable and cost-effective approach to measure broad antibody binding specificity and to predict specific off-target interactions does not exist. This lack of a broad and predictive measure of specificity limits the ability to identify antibody clones with a desired specificity profile at the early stages of discovery.

Immunohistochemistry and Western Blots are methods currently used to determine binding specificity for a small number of selected clones, but are not scalable to large panels of antibody candidates identified in a screen.^63–66^ Protein microarrays can be used to assess broad antibody binding specificity,^67–69^ but the high cost of these arrays limits specificity measurements that would ideally be performed in the early stages with hundreds of selected clones. Also, protein microarrays do not provide sequence-level information about the off-target interaction(s) that would be required for prediction of off-target binding partners. Antibodies are generally believed to have exquisite binding specificity, but it is important to identify any off-target interactions early because several examples exist where off-target interactions severely impacted therapeutic antibody efficacy.^61, 62, 70^

An array-based peptide library platform for high-resolution antibody binding characterization produced at the scale, quality and assay format required for early-stage antibody discovery is described here. This platform aims to close the gap between the large numbers of potentially valuable clones that can be identified in a screen and the limited number of clones that can be characterized. The current antibody characterization paradigm involves an initial epitope binning step followed by epitope mapping on a few clones that are representative of the epitope bins. The platform described in this study enables epitope mapping for each clone that needs to be characterized, leading to subsequent binning of all clones based on their array binding characteristics. This approach is a fundamental shift in the epitope characterization process when compared to the current approach to epitope binning and mapping that is highly iterative. In addition, epitope mapping and binding specificity can be performed in parallel with this platform, early in the antibody screening workflow. Importantly, the information content generated from array-based peptide library antibody binding is sufficient to predict potential off-target interaction partners for an antibody clone. This off-target prediction capability is a new early-stage antibody discovery characterization metric enabled by this platform.

To achieve these advancements in antibody epitope and specificity characterization, array-based peptide library synthesis has been developed using photolithographic masks (ref. 71) designed to produce 3.3M peptides that can cover a full hexamer sequence space. This 3.3M peptide library is not designed using prior target or epitope knowledge which enables *de novo* epitope identification on a target. Competition-free epitope binning and putative epitope mapping is done by aligning the top antibody binding peptides from the 3.3M peptide library to the target protein. The full-hexamer space binding results are then compared to antibody binding on a 126K peptide diverse sequence space library. High-resolution epitope characterization is achieved on this platform with array-based 17K peptide focused sequence space peptide libraries to measure position-level binding contributions and selectivity. These array-based 17K peptide focused libraries are analogous to site-directed and shotgun mutagenesis libraries. Finally, broad and predictive off-target antibody binding specificity characterization is performed using antibody binding results from both the 3.3M full and 126K sequence space libraries.

## Results

A panel of anti-HER2 monoclonal antibodies (mAbs) known to bind to different regions of HER2 (Table 1) were used to demonstrate epitope binning, mapping, and specificity characterization with full, diverse, and focused sequence space array-based library designs. Each of these monoclonal antibodies have known immunogens that were used as a comparative reference for the array-based epitope binning and mapping results.

**Table 1:** Anti-HER2 mAb panel and immunogens.

| Clone | Host | Immunogen<br>Region |
| --- | --- | --- |
| D8F12 | Rabbit | N-terminus |
| C-3 | Mouse | 251-450 |
| C-12 | Mouse | 983-1017 |
| A-2 | Mouse | 1180-1197 |
| EP1045Y | Rabbit | 1200-1255 |
| 9G6 | Mouse | 1200-1255 |
| SPM495 | Mouse | 1200-1255 |
| F-11 | Mouse | 1238-1255 |
| SC-3B5 | Mouse | 1242-1255 |
| TF-3B5 | Mouse | 1242-1255 |
| 44E7 | Mouse | 1242-1255 |
| 29D8 | Rabbit | 1242-1255 |
| RM228 | Rabbit | 1242-1255 |
| QO3B | Mouse | 1242-1255 |
| A24-V | Rabbit | 1242-1255 |

Two array-based broad peptide sequence space library design approaches were evaluated for epitope binning and mapping. First, a 16-well plate format (one array/well) full coverage library was synthesized that includes 3M hexamers and 125K decamers using a reduced 12-amino alphabet, {G, V, P, L, W, Y, S, N, K, R, H, D}, derived from BLOSUM amino acid similarity matrices.^72^ In this full coverage library, all possible 3M 12-amino acid alphabet hexamers are included along with 125K decamers designed to maximize heptamer diversity within the context of decamers (500K distinct heptamers within the context of 125K decamers). This design generates a total of 3.3M array peptides that includes library replicates and control sequences. A second 96-well plate format (one array/well) diverse coverage library of 126K peptides was synthesized using a 16-amino acid alphabet that excludes oxidation-prone cysteine/methionine residues and utilizes amino acid similarity by substituting threonine and isoleucine with serine and valine respectively. This diverse sequence space library was designed to have a median peptide length of 9 with 100% of tetramer space and 50% of pentamer space coverage.^73^ Both of these broad sequence space library designs do not utilize knowledge of the HER2 immunogens, thereby enabling epitope binning, mapping, and specificity determination across a wide range of antibody interactions.

A third array-based 96-well plate format (one array/well) focused peptide sequence space library containing 17K sequences was evaluated for high resolution epitope mapping. Focused sequence space libraries were synthesized photolithographically using a physical mask approach that defines a fraction of overlapping holes between sequential masks in the full mask series. The peptide library designer can then introduce “mutations” at defined peptide library positions with a desired percentage occurrence by selecting sequential masks with that same fraction of overlapping holes, analogous to designing a site-directed mutagenesis library. An inherent characteristic of the sequential overlap mask approach is a statistical population of overlapping holes on two or more non-sequential masks that are analogous to random mutagenesis libraries. Taken together, this sequential overlap mask approach produces an array-based library of 17K peptide sequences that includes cognate sequences, point mutants, multi-position mutants, deletions, truncations and divergent sequences.

For all library designs, antibody binding assays were run in an immunogen competition format where the anti-HER2 monoclonal antibody panel was pre-incubated with (+/-)HER2 overexpression cell lysate competitor, then bound to the peptide arrays. HER2 expressing lysate competitor was chosen for two reasons, first to present both HER2 intracellular and extracellular native domains as competitor to the mAbs in the panel, and second to demonstrate platform compatibility of antibody binding within a complex biological matrix. This antibody-target competition assay format is different in design and intention when compared to pairwise antibody-antibody cross-blocking competition assays used for epitope binning. In this case, the number of measurements does not increase exponentially with the number of clones to be characterized and does not intend to identify cross-blocking bins. Instead this antibody-target competition assay format intends to improve the target epitope relevance of peptide array antibody binding results in two ways. First, this assay format provides an indirect measure of relative peptide-antibody binding energy and the effect of positional substitutions for all peptides on the array, with respect to the cognate antibody-target interaction in solution. Second, this assay format confers a level of specificity in the peptide array binding results since, in the presence of native immunogen, only those peptide array interactions most reflective of the cognate antibody-target interaction will retain their binding signal.

### In Situ MALDI-MS characterization of photolithographic peptide array synthesis

Comprehensive *in situ* MALDI-MS capabilities are built into the peptide array platform to ensure that peptides on the array are efficiently synthesized. Importantly, MALDI-MS measurements are taken *in situ* from the same silicon wafer that contains the synthesized peptide array libraries. This approach ensures MALDI-MS synthesis characterization reflects library quality at a per-synthesis level and is not extrapolated across many syntheses. An overlapping trimer and hexamer approach was built into the peptide array synthesis platform to cover all chemical steps required to synthesize up to 3.3M peptides and to calculate synthesis cycle efficiency. *In situ* MALDI-MS spectra from overlapping hexamers and trimers (Figs. 1, S1) indicate typical synthesis cycle efficiencies greater than 98% and peptide purity greater than 90% (Tables 2, S1).

**Figure 1:**
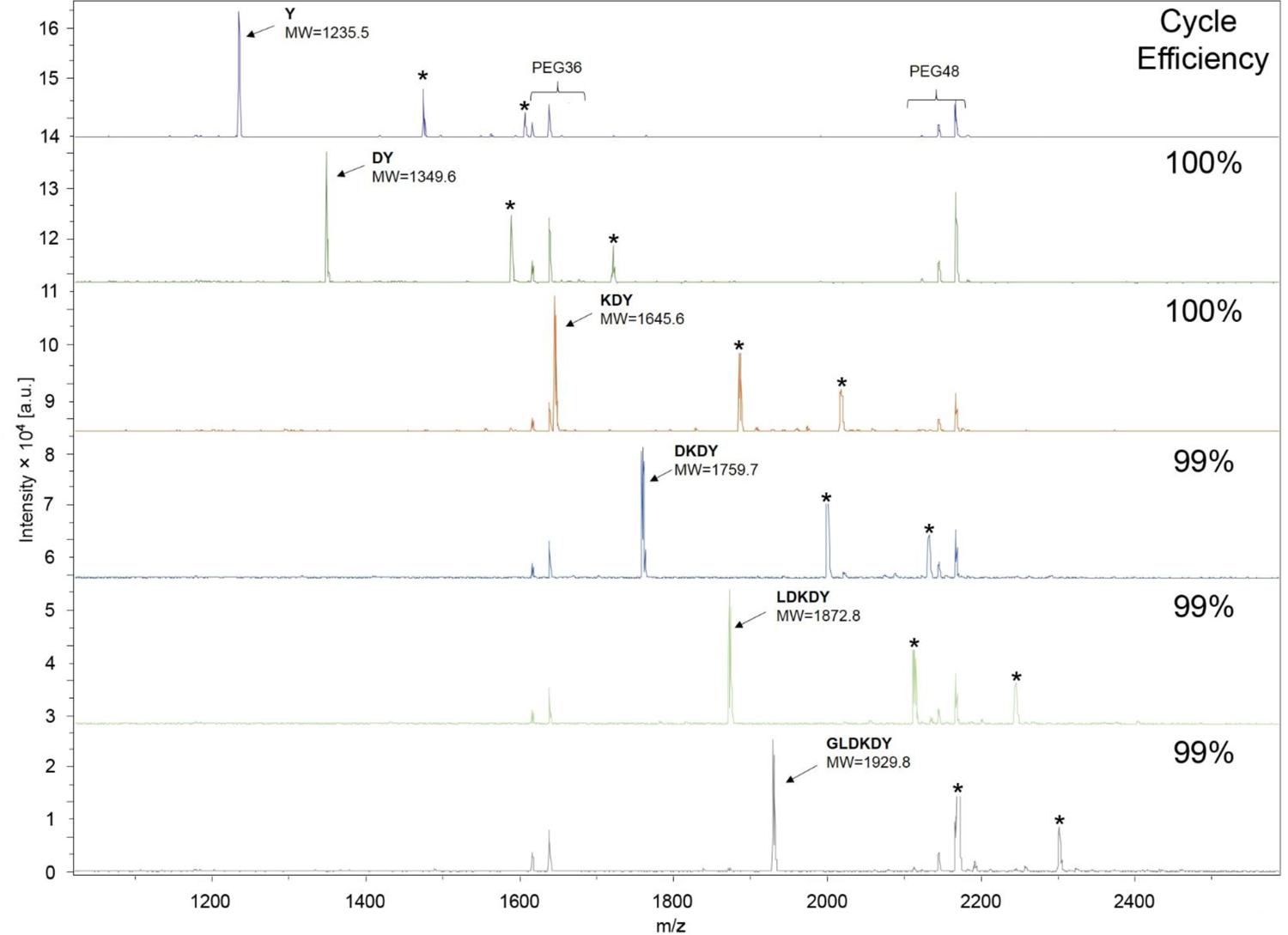
*In situ* MALDI-MS length series of photolithographic hexamer synthesis. Peptide length series are used to verify the successful consecutive photo-deprotection and coupling of amino acids. A stacked 1-6 peptide length series of six MALDI spectra leading up to the GLDKDY hexamer are shown. Each mass spectrum is acquired *in situ* from distinct array features corresponding to the peptide length 1-6. The expected m/z peak for each length product are marked by arrows and correspond to the peptide product plus an N-terminal tris (2,4,6-trimethoxyphenyl)phosphonium-acetyl (TMPP-Ac) group and a C-terminal 6-residue polyethylene glycol (PEG) linker. Two mono-dispersed PEG-amines, PEG36 and PEG48 were added in the MALDI matrix as standards for calibration (PEG [M+H] and [M+Na] peaks observed). (*) Known alternate linker cleavage products that include desired peptide: [M+240], [M+371].

**Table 2:** *In Situ* MALDI-MS characterization of photolithographic synthesis cycle efficiency. Each product mass spectrum is acquired *in situ* from a distinct array spot. Product purity and cycle efficiency are calculated from the product length (n = 1 - 6) series of spectra. Cycle efficiency for product length n is relative to the n - 1 length product. Cycle efficiency for the n = 1 length is derived from mass spectrum substitution peaks that could arise from subsequent amino acid couplings if the first cycle did not yield 100% expected product.

| Cycle<br>(n) | Expected<br>Product | Mass-to-Charge | Signal-to-Noise | Product<br>Purity | Cycle<br>Efficiency |
| --- | --- | --- | --- | --- | --- |
|  |  | (m/z) | (S/N) | % (+/- 2) | % (+/- 2) |
| 1 | Y | 1235.5 | 329 | 100 | 100 |
| 2 | DY | 1349.6 | 156 | 100 | 100 |
| 3 | KDY | 1645.6 | 414 | 99 | 100 |
| 4 | DKDY | 1759.7 | 335 | 96 | 99 |
| 5 | LDKDY | 1872.8 | 298 | 96 | 99 |
| 6 | GLDKDY | 1929.8 | 268 | 93 | 99 |

### Monoclonal antibody epitope binning and putative epitope mapping with an array-based ***3.3M full-coverage proteome-representative hexamer library***

Full coverage of 3M hexamer sequence space for epitope binning and mapping was achieved with a 3.3M feature peptide array (7 μm feature diameter, 11 μm pitch), with one array feature corresponding to one peptide sequence. Full hexamer coverage is attained using a reduced 12-amino acid alphabet where the key design principle behind the reduced alphabet is BLOSUM amino acid similarity (Fig. S2).^72^ By using BLOSUM amino acid similarity derived from substitution degeneracy observed in the proteome, the 3.3M feature array is compositionally representative of the much larger 20 amino acid alphabet proteome sequence space. Also, by covering the full sequence space, this library expands beyond annotated hexamer sequence coverage in the proteome. Important for epitope binning and mapping, antibody binding assays performed with this proteome representative library produce sequence hit lists that can be aligned to the immunogen sequence using preferred sequence alignment algorithm(s). These putative epitope alignments can be scored using the same BLOSUM similarity matrix used to design the library.^74^ A reduced 12-amino acid alphabet 3.3M peptide array library does have potential shortcomings in cases where particular antibody clones have higher selectivity and/or affinity for one BLOSUM similar amino acid vs. the alternate, for example glycine vs. alanine. However, coverage of all possible sequences with a reduced alphabet offers the advantage of sequence-level antibody characterization vs. all possible sequences in the space without the risk of missing multi-position dependencies that would not be represented in peptide libraries without full sequence space coverage. If needed, fully resolved amino acid binding selectivity of antibodies can be measured with focused peptide array libraries that include the full amino acid alphabet.

Peptide consensus logos were generated for 15 anti-HER2 mAbs using the top five binding peptides from the 3.3M peptide array in the presence of HER2 expressing cell lysate. For each clone, putative epitope bins were identified by gapped pairwise alignment of the conserved top peptide motifs to the HER2 sequence (Fig. 2). Binning antibodies based on this direct array binding approach is significantly more scalable than traditional pairwise mAb cross-blocking epitope binning, where typically for N number of clones at least N^2^/2 pairwise measurements are required, compared to just N measurements with this approach. Array binding produced highly conserved top peptide sequence motifs for clones 29D8, TF-3B5, SC-3B5, QO3B, RM228, A24-V, SPM495, EP105Y, F-11 and C-3 that allowed epitope binning by alignment of the observed consensus motifs to the full HER2 primary sequence. Lower sequence conservation in top peptide motifs was observed with clones D8F12, C-12, A-2, 44E7 and 9G6. In these cases, motif pairwise alignment for epitope binning required restriction to HER2 regions surrounding the immunogen. This observed difference in the level of top peptide motif conservation for different clones may be the result of varying per-clone binding specificity.

**Figure 2:**
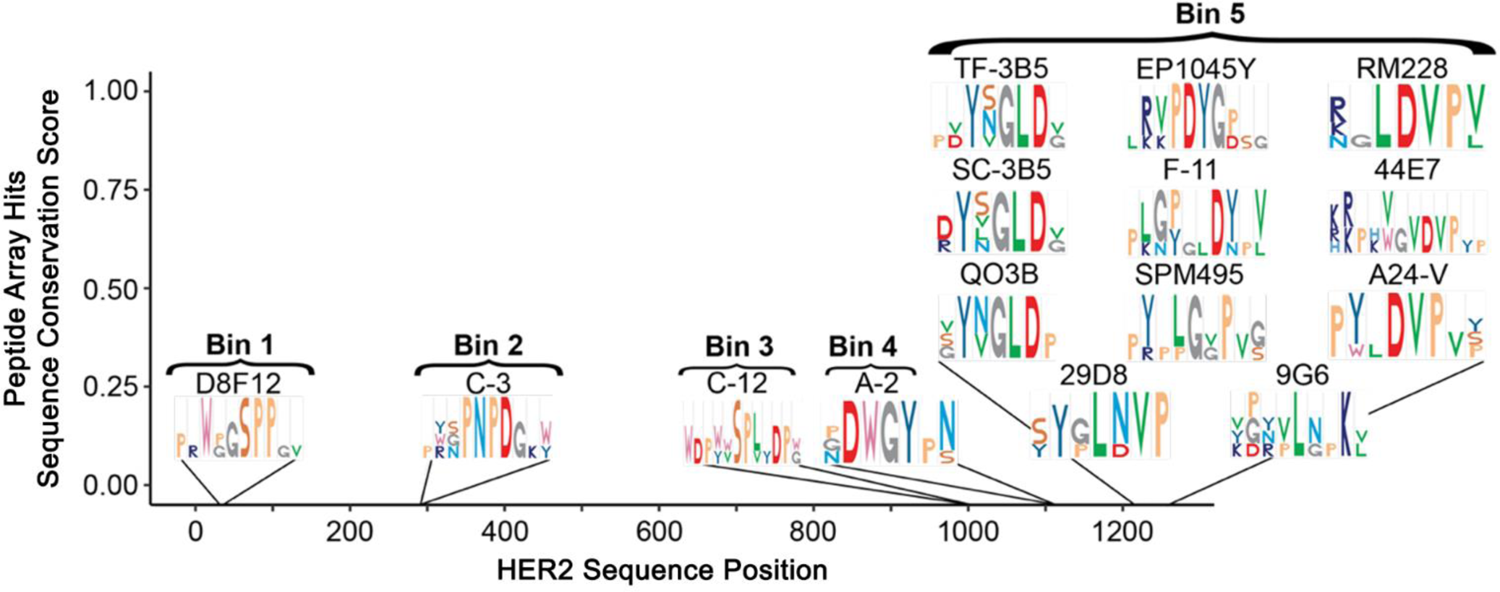
Putative HER2 epitope bins were identified using the 3.3M full-hexamer coverage library top five binding peptides. Top peptide position-level sequence conservation scores (y-axis) were generated for each anti-HER2 clone using Eq. 1 in the methods section. Scored conserved motifs, shown as Weblogos, were pairwise aligned to the HER2 primary sequence (x-axis) to generate putative epitope bins.

Putative HER2 epitope mapping was performed using the same top five mAb binding peptides from the 3.3M peptide array as used for epitope binning. Utilizing the top five peptide hits vs. more than five hits for epitope mapping was a balance between binding information content and sequence alignment performance. A gapped multiple-sequence alignment strategy was used to map clones to putative epitopes with the top peptides aligned to the HER2 immunogen sequence (Fig. 3). These alignments are scored using the same BLOSUM similarity matrix used to produce the reduced 12-amino acid alphabet for the library design. With multiple-sequence alignment and BLOSUM scoring, a target coverage score can be generated for each aligned position to highlight critical binding residues in the putative epitope. Typically, across the 15 anti-HER2 mAb panel, 4-5 key conserved residues in a parent 6-mer peptide sequence were observed in the top binders (Fig. 3). In addition, since the hexamer peptide library has full coverage of the sequence space, stacked and shifted alignment of several peptides provides more than six coverage positions in the target sequence. Highlighting reproducibility, the anti-HER2 3B5 clone from two different vendors (TF-3B5 and SC-3B5) overlap in three of five top binding peptides, out of 3.3M total peptides, and their target coverage is nearly identical.

**Figure 3:**
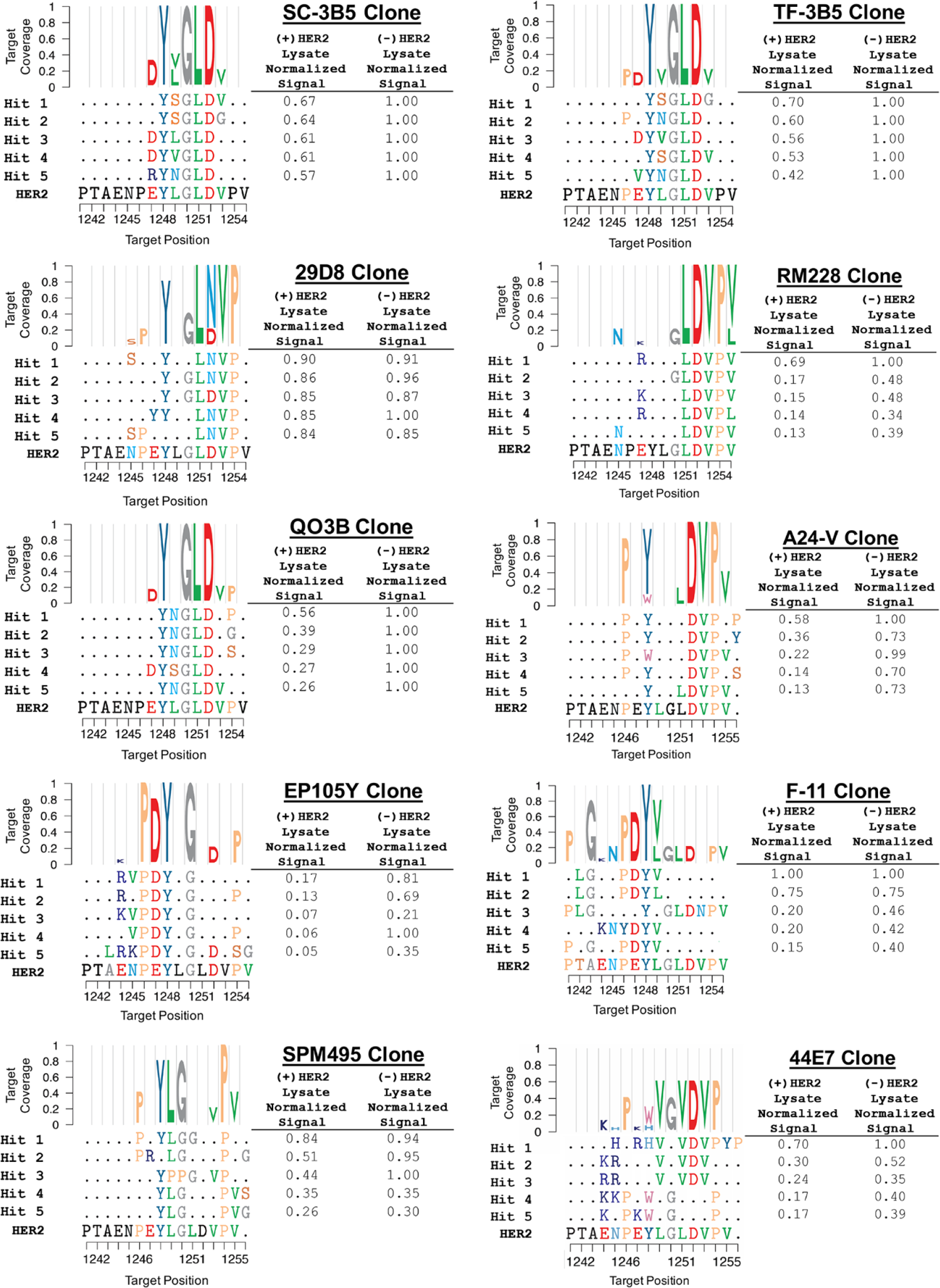
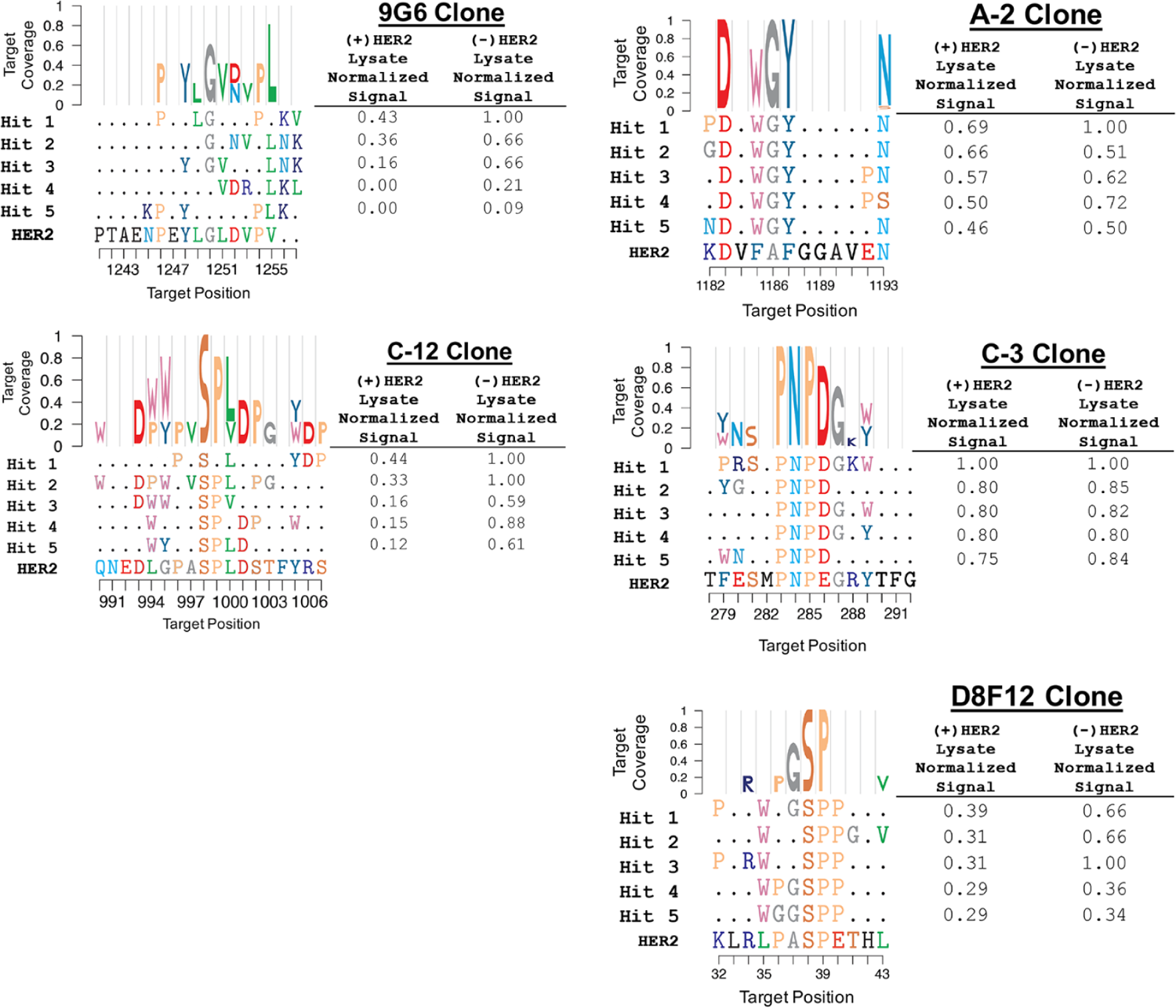
Putative epitope mapping using the top five 3.3M full-hexamer coverage library binding peptides for each clone. Top peptides for each clone aligned to the Her2 sequence (x-axis) are shown. Each aligned position is scored for target coverage (y-axis) using Eq. 2 in the methods section and those scores are used to generate Weblogos. Relative array binding signal for each top peptide for the (+/-)HER2 competitor lysate conditions is shown in the table to the right of each alignment and logo.

Antibody panel top binding peptide results on the 3.3M peptide array showed a range of putative HER2 epitope coverage that may be indicative of a range of antibody clone binding affinity and specificity characteristics. Like the conserved motifs used for epitope binning, high scoring top peptide putative epitope mapping coverage was observed with clones 29D8, TF-3B5, SC-3B5, QO3B, RM228, A24-V, SPM495, EP105Y, F-11 and C-3. In these cases, epitope mapping could be performed with multiple sequence alignment to the full HER2 primary sequence. Reflective of epitope binning conserved top peptide motifs, lower scoring epitope coverage was observed with clones D8F12, C-12, A-2, 44E7 and 9G6. In these cases, putative epitope mapping required limiting the alignment of top array binding sequences to the HER2 primary sequence regions surrounding the known HER2 immunogen. Finally, significant variation in tolerance to amino acid substitutions relative to the HER2 cognate sequence was observed across the anti-HER2 mAb panel. Particularly striking in the clones immunized with a C-terminal HER2 immunogen is the tolerance to the D1252N substitution for clone 29D8 but not clones TF-3B5, SC-3B5, QO3B or RM228. The ability to measure this tolerance to substitution in high-resolution could identify antibody clones with cross-species reactivity or even tolerance to position-specific tumor mutations.

### Monoclonal antibody binding to an array-based 126K diverse 16-amino acid alphabet sequence space library

Ten anti-HER2 mAb clones using an identical (+/-)HER2 HEK293 cell lysate assay format were bound to an array-based 126K diverse sequence space peptide library (14 μm feature diameter, 18 μm pitch). This library was synthesized with a 16-amino acid alphabet to compare epitope mapping using a diverse, but not full-coverage, sequence space library to the 3.3M 12-amino acid alphabet full-hexamer coverage sequence space epitope mapping results. The diverse-coverage library contains over 500K distinct hexamer sequences within the median 9-mer context. In addition to comparing epitope mapping resolution between full-coverage and diverse-coverage libraries, this comparison was performed to gain insight on future peptide array library designs that could improve antibody characterization information content, throughput or cost.

To maintain putative epitope mapping analysis consistency, the top five antibody binding peptides from the diverse-coverage library were aligned to the known immunogen region using gapped multiple sequence alignment and scored using the same BLOSUM similarity matrix used for the 3.3M full-coverage library alignments (Fig. 4). One notable difference in epitope mapping alignment using diverse peptide sequence libraries is that the non-cognate residues flanking the cognate epitope residue matches reduced alignment performance when standard multiple sequence alignment algorithms such as Clustal Omega or Muscle where used.^75, 76^ To address this, epitope mapping alignments of top binding diverse library sequences were limited to HER2 sequence regions surrounding the known immunogen.

**Figure 4:**
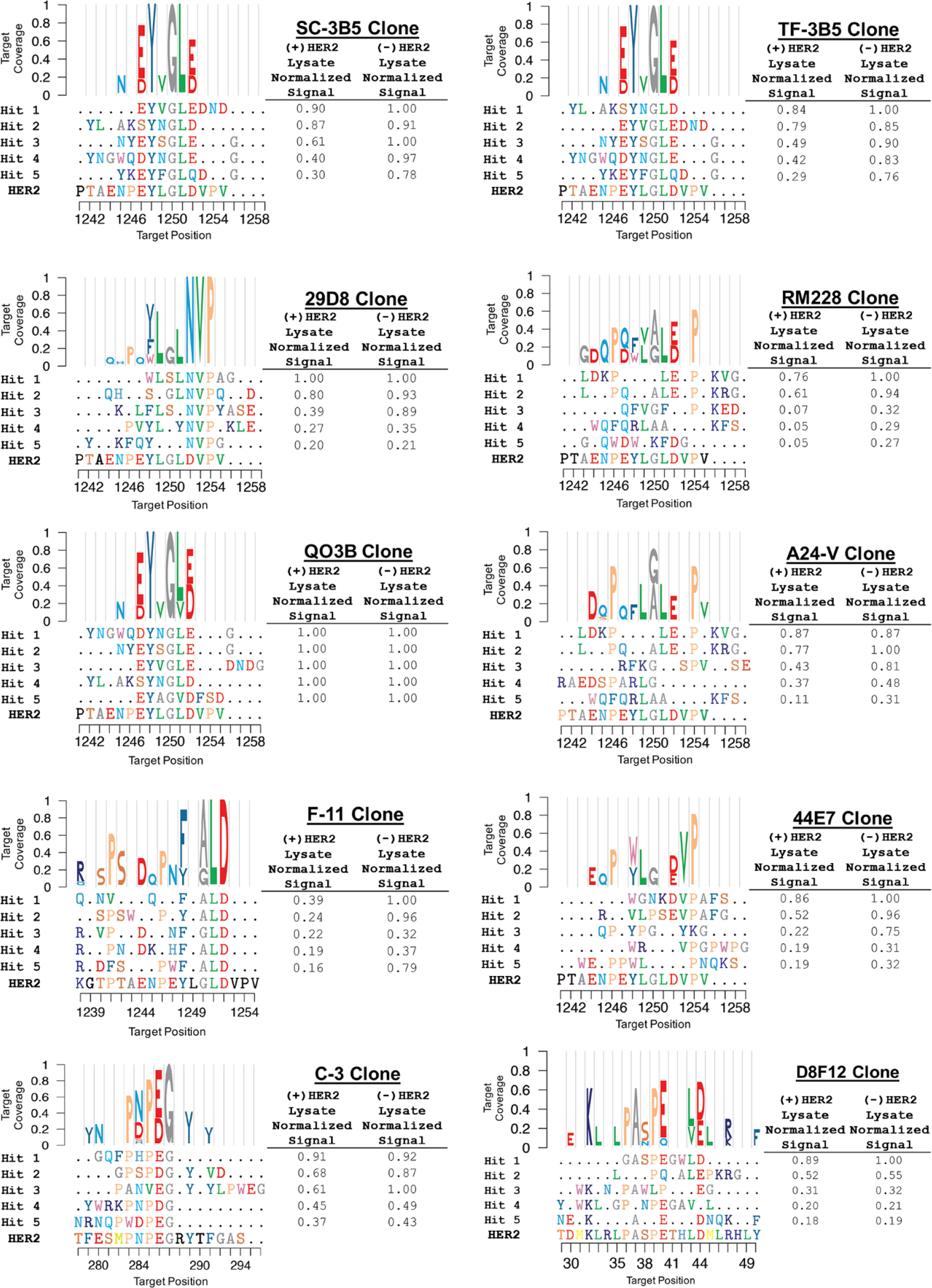
Putative epitope mapping using the top five 126K diverse coverage library binding peptides for each clone. Top peptides for each clone aligned to the HER2 sequence (x-axis) are shown. Each aligned position is scored for target coverage (y-axis) using Eq. 2 in the methods section and those scores are used to generate Weblogos. Relative array binding signal for each top peptide for the (+/-)HER2 competitor lysate conditions is shown in the table to the right of each alignment and logo.

Conserved motif and target coverage are in strong agreement between diverse and full-coverage libraries for clones 29D8, TF-3B5, SC-3B5, QO3B and C-3. As was observed with the 3.3M full-coverage library, diverse library top binding peptide alignments highlight differences in substitution tolerance between clones. The same noteworthy example is observed in the diverse library data where clone 29D8 tolerates the D1252N substitution, but clones TF-3B5, SC-3B5, QO3B or RM228 do not. The top peptide hit from the diverse sequence library for clone D8F12, GASPEGWLD, is an interesting case that may be informative on improving library design for epitope mapping with peptide arrays. In this case, the diverse library D8F12 clone top peptide hit, GASPEGWLD, has a high number of alignment position matches with the known HER2 immunogen relative to the total length of the peptide. Also, this peptide hit retains most of its array binding signal in the presence of (+)HER2 lysate competitor. Similar retention of binding signal in the (+)HER2 competitor condition is observed in other top diverse library binding peptides that have a high number of alignment position matches with their known immunogen. This result suggests peptide libraries that maximize the number of alignment matches relative to the peptide length could improve epitope mapping and critical residue identification and this is one motivation behind the focused library design described in the following section. As was observed with 3.3M full-coverage library binding, low scoring target coverage and consensus was observed on diverse-coverage library binding with clones 44E7 and D8F12. This may suggest that these clones are more promiscuous in binding peptide/protein sequence space relative to the other clones tested. Diverse and full-coverage sequence space binding for clone F-11 produced target coverage and consensus corresponding to the known immunogen in both cases, but the target coverage and consensus are different for the two library designs. The top F-11 clone binding peptides from the diverse peptide library have high FALD motif conservation, whereas the top peptides from the 3.3M full sequence space library have high LGPDYV motif conservation. Each of these motif conservations are compositionally related to the cognate HER2 immunogen sequence. This may indicate that the paratope of the F-11 clone can accommodate different orientations of the key energetic residues, one orientation with the Y/F residue N-terminal and another with Y/F C-terminal. Like the top peptide hit reproducibility observed with the 3.3M full hexamer-coverage library, the single clone 3B5 acquired from two different vendors (TF-3B5 and SC-3B5) produce identical top five binding peptide hits using the 126K diverse library with a slightly different rank-order (Fig. 4).

### High-Resolution epitope mapping with an array-based 17K focused-coverage sequence space library

Two focused libraries were synthesized for high-resolution epitope characterization. These libraries utilized a recently developed photolithographic mask strategy described above to produce an array-based library of 17K peptides (44 μm feature diameter, 50 μm pitch) focused on putative HER2 epitopes. Specifically, these two libraries were focused on putative epitope regions 1242-1255 and 275-295, identified in the 3.3M full-coverage library epitope mapping results (Fig. 3). Focused libraries included cognate sequences, point and multiple substitution mutants, deletions and truncations derived from the putative epitope regions along with divergent sequences to identify potential antibody off-target binding interactions. This focused library design corroborates and improves the resolution of the diverse libraries in the following ways: 1) Provide confirmation of putative epitopes identified on 3.3M full-hexamer libraries, 2) Characterize position-level epitope differences between clones, 3) Identify key energetic epitope binding residues, and 4) Characterize residue-level tolerance to substitutions, deletions and truncations between clones.

Putative epitope mapping using the 3.3M full-coverage library identified HER2 residues 1242-1255 as a putative epitope for antibody clones: 29D8, TF-3B5, SC-3B5, QO3B, RM228, A24-V, 44E7, F-11, and residues 275-295 as a putative epitope for clone C-3 (Fig. 3). These clones were bound to their corresponding focused library arrays. Alignments of top binding peptides identified from the focused-coverage libraries were done using the same sequence alignment and BLOSUM similarity scoring matrix approach used for the 3.3M full-coverage and 126K diverse-coverage library alignments. High sequence consensus and putative epitope coverage were observed in the top binding sequences for all clones assayed on their respective focused peptide library (Fig. 5), even though most of the 17K focused peptide library did not include the observed conserved motifs. This suggests that the observed binding motifs are not driven by sequence biases in the focused library. Focused library sequence conservation and epitope coverage scores corroborate the 3.3M full-coverage library as well as 126K diverse-library top peptide results (Figs. 3, 4). In all but two cases (clones RM228, 44E7), the top focused-coverage library binding peptides identified for each clone compete at near equivalence in the presence of (+)HER2 overexpression cell lysate relative to the (-)HER2 lysate. Taken together, high sequence consensus, epitope coverage, and (+/-)HER2 competition equivalence, these data suggest that the top focused-library binding peptides can identify most or all of the key energetic epitope residues for a particular clone.

**Figure 5:**
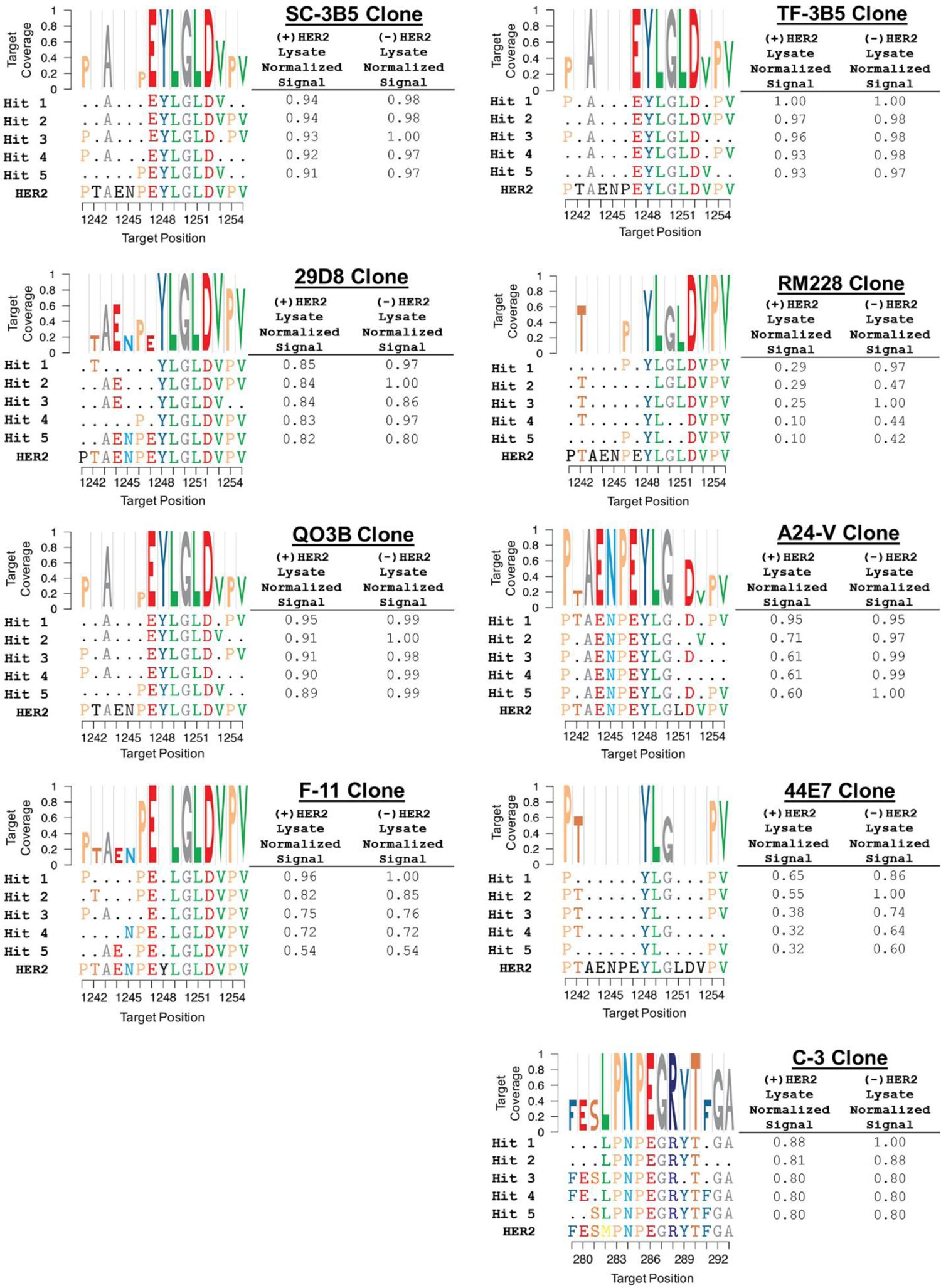
High-resolution epitope mapping using the top five 17K focused coverage library binding peptides for each clone. Top peptides for each clone aligned to the HER2 sequence (x-axis) are shown. Each aligned position is scored for target coverage (y-axis) using Eq. 2 in the methods section and those scores are used to generate Weblogos. Relative array binding signal for each top peptide for the (+/-)HER2 competitor lysate conditions is shown in the table to the right of each alignment and logo.

Even with the relatively short 1242-1255 HER2 immunogen used to raise clones 29D8, TF-3B5, SC-3B5, QO3B, RM228, A24-V, 44E7 and F-11, significant position-level epitope differences are observed (Fig. 5). For example, many C-terminal HER2 immunogen clones have their energetic epitope residues near the C-terminus of the 1242-1255 immunogen, but with this high-resolution epitope mapping data, clone A24-V would fall into a distinct epitope bin towards the N-terminus of this immunogen (Fig. 5). High epitope mapping reproducibility is observed with the single 3B5 clone acquired from two vendors using the 17K focused-library (Fig. 5: TF-3B5, SC-3B5), as was seen in the 3.3M full-coverage and the 126K diverse-coverage libraries.

### Antibody specificity characterization and off-target prediction

To address the need for a scalable, quantitative approach that reflects broad antibody binding specificity, top array binding peptide epitope coverage scores were evaluated as a measure of specificity. For each clone bound to both full hexamer coverage and diverse sequence space arrays, an overall coverage metric was calculated as a sum of all aligned top peptide array hit positions with a coverage score >0.50 and normalized to the total number of aligned positions in the peptide hit set. The coverage metric for each clone was then scaled to 1 across the full panel, between both full hexamer and diverse sequence space libraries, and rank-ordered (Table 3). The hypothesis is that a higher ranked coverage metric would reflect higher HER2 binding specificity in a broad protein background. To test this hypothesis, each clone was used in a Western Blot to probe binding in (+/-)HER2 overexpressing HEK293T cell lysates. In the Western Blots, the number of bands present in addition to the HER2 band serves as an orthogonal measurement of HER2 antibody binding specificity in a broad protein background. In general, anti-HER2 clones with a lower epitope coverage metric calculated from the top peptide array binding sequences have more off-target binding bands in the cell lysate Western Blot (Fig. S3). This is particularly the case when binding to both 3.3M full-coverage and 126K diverse-coverage libraries is considered.

**Table 3:** Epitope coverage metrics for top binding peptides from full-hexamer coverage and diverse sequence space libraries. An epitope coverage metric was calculated using the top five array binding peptides from the 3.3M full-hexamer and the 126K diverse coverage libraries and (Eq. 3) described in the methods. Each metric is included in the corresponding clone Western Blot (Fig. S3) as a correlate to broad binding specificity in a cell lysate background.

|  | 3.3M Full-Hexamer Library | 126K Diverse Library | Full & Diverse Libraries |
| --- | --- | --- | --- |
| Clone | Epitope Coverage Metric | Epitope Coverage Metric | Epitope Coverage Metric (Mean) |
| SC-3B5 | 1.00 | 0.51 | 0.76 |
| TF-3B5 | 0.88 | 0.51 | 0.70 |
| QO3B | 0.88 | 0.45 | 0.67 |
| 29D8 | 0.84 | 0.44 | 0.64 |
| C-3 | 0.70 | 0.36 | 0.53 |
| RM228 | 0.80 | 0.19 | 0.50 |
| F-11 | 0.45 | 0.53 | 0.49 |
| A24-V | 0.75 | 0.18 | 0.47 |
| D8F12 | 0.38 | 0.29 | 0.34 |
| 44E7 | 0.40 | 0.20 | 0.30 |

In addition to broad binding specificity, top binding peptides and/or highly conserved motifs from 3.3M full-coverage sequence space binding assays may be predictive of off-target binding partners for an antibody. In these putative off-target cases, the conserved array binding motifs do not exactly match the on-target epitope sequence, but instead match off-target sequence(s) identified in a proteome search. Potential off-target antibody binding interactions were predicted for the three clones that exhibited the highest HER2 binding specificity, based on peptide array binding epitope coverage metrics (Table 3) and (+/-)HER2 expressing cell lysate Western Blots (Fig. S3). Putative antibody off-target interaction sequences were identified as those top peptide binding sequences with non-cognate substitutions and retention of 50% their relative signal in the (+)HER2 expression lysate array binding conditions (Fig. 3). These non-cognate substituted off-target sequences are predicted to have affinity comparable to the cognate epitope interaction based on (+/-)HER2 lysate array binding assays. Putative off-target sequence motifs were searched against the proteome database for potential off-target binding partners. Affinity for the YSGLD peptide motif was identified for clone 3B5 from both vendors (TF-3B5, SC-3B5) and clone QO3B but not clone C-3, which served as a negative control for dot-blot assays. A proteome search for the YSGLD motif predicted ATF-6 Beta (Uniprot: Q99941) as an off-target binding partner (ATF-6 Beta region 35-39). Dot-blot binding assays using purified recombinant ATF-6 Beta with clones TF-3B5, SC-3B5, QO3B, and C-3 corroborate array binding off-target predictions. Clones TF-3B5, SC-3B5, QO3B bind ATF-6 Beta at 0.4 and 0.04 μg/mL doses at an intensity comparable to (+)HER2 expressing cell lysate (Table 4) but clone C-3 does not bind ATF-6 beta. Clone TF-3B5 array binding results identified the sequence PYNGLD as a potential specific off-target binding interaction with Her4 (Uniprot: Q15303) that could compete with HER2 binding, where HER4 contains a highly similar PYNAIE motif at region 377-382. Again, a dot-blot binding assay corroborates array-predicted off-target binding where HER4 binding is observed with clone TF-3B5, but not clones SC-3B5, QO3B, and C-3 (Fig. S4). Since TF-3B5 and SC-3B5 are technically the same clone, but from two vendors, this off-target result for TF-3B5 may suggest a production and/or lot difference in TF-3B5 vs. SC-3B5.

**Table 4:** Array-predicted off-target binding intensities observed in a dot blot assay. Dot-blot binding intensities (dot-blot image, Fig. S4) for the three highest scoring epitope coverage metric mAbs, SC-3B5, TF-3B5, and QO3B along with the median scoring mAb C-3 were probed for array-predicted off-target binding. Using the 3.3M full-hexamer library top five array binding sequences, clone TF-3B5 was predicted to bind HER4 and mAbs SC-3B5, TF-3B5 and QO3B were predicted to bind ATF-6 Beta. Clone C-3 served as a negative control for these predicted off-target interactions. HER2 dot-blot binding intensities are highlighted in green, predicted off-target binding intensities are highlighted in red.

| Clone | C-3 |  | SC-3B5 |  | TF-3B5 |  | QO3B |  |
| --- | --- | --- | --- | --- | --- | --- | --- | --- |
|  | Background Subtracted Intensity |  | Background Subtracted Intensity |  | Background Subtracted Intensity |  | Background Subtracted Intensity |  |
|  | 0.4 | 0.04 | 0.4 | 0.04 | 0.4 | 0.04 | 0.4 | 0.04 |
| Protein | mg/mL | mg/mL | mg/mL | mg/mL | mg/mL | mg/mL | mg/mL | mg/mL |
| HER4 | 0 | 0 | 488 | 0 | 4658 | 252 | 1824 | 0 |
| ATF-6 Beta | 198 | 882 | 11156 | 3024 | 16387 | 1400 | 4516 | 306 |
| (+) HER2 Lysate | 4246 | 228 | 10854 | 7708 | 12688 | 7388 | 9190 | 324 |
| (-) HER2 Lysate | 192 | 0 | 0 | 264 | 0 | 0 | 274 | 0 |

## Discussion

Comprehensive epitope identification and characterization is a key part of early-stage therapeutic antibody discovery because an antibody’s epitope is a primary determinant of efficacy.^14^ Two important aspects of epitope characterization include: 1) binning antibody clones based on their ability to compete for an epitope,^22–25, 30–32^ and 2) high-resolution epitope mapping to identify the target binding site and critical residues involved in binding (i.e. the functional epitope).^33–49^ The traditional cross-blocking epitope binning assay format requires at least N^2^/2 pairwise measurements for N clones.^22–25, 30–32^ High-resolution epitope mapping approaches produce a comprehensive picture of the antibody-target interaction but are resource-intensive.^33–, 49^ Taken together, the high number of epitope binning measurements and the resource requirements of epitope mapping severely limit the number of clones that can be characterized in early-stage antibody discovery relative to the large number of clones identified in a screen.

Characterization of antibody binding specificity is another important aspect of antibody discovery. Off-target antibody interactions arising through the CDR-loops or any other region of an antibody can produce side-effects that impact efficacy.^61, 62, 70^ Identifying these off-target interactions early in antibody discovery could reduce the risk of side-effects observed in vivo. The current assay platforms available for antibody specificity determination are limited in both throughput and information content.^63–69^ Current approaches not only limit specificity characterization to a small number of antibody clones, but also do not provide sufficient information to identify the motif(s) responsible for the off-target interaction(s) that could then be used to predict potential off-target binding partner(s).

Filling the existing gap in antibody discovery between the large numbers of antibodies selected in a screen and the limited numbers of antibodies that can be characterized could significantly improve antibody discovery productivity. Also, a platform that provides the capability to characterize hundreds of antibody clones in the early stages of discovery would increase the likelihood of identifying clones with exquisite specificity or other novel binding characteristics not easily identified with current characterization platforms. *In situ* synthesized peptide microarrays are one attractive platform to fill the high-throughput antibody characterization gap due to the ability to pattern these libraries in high density, their high sequence information content, and their compatibility with complex sample matrices.^52–54, 58^ However, the application of *in situ* synthesized peptide microarrays in early-stage antibody discovery has not been fully realized to-date due to limited array production scalability, high-throughput automation incompatibility, and a lack of comprehensive per-synthesis quality assessment.^52, 59, 60^

We have developed a scalable, high-quality peptide array platform to address the need for increased characterization throughput and information content in early-stage antibody discovery. This platform was developed with sequence space coverage flexibility for various levels of resolution in epitope and specificity characterization. Epitope binning, mapping and specificity characterization was demonstrated using this platform and a panel of anti-HER2 monoclonal antibodies with three array-based peptide library design strategies that deliver different degrees of information content. This study included a 3.3M peptide 12-amino acid alphabet full-coverage library design, a 126K peptide 16-amino acid alphabet diverse-coverage library, and a 17K peptide complete amino acid alphabet focused-coverage library. These libraries are produced in a format for high-throughput automation with one array per well (Fig. S5). Notably, using the photolithographic synthesis conditions described here, comprehensive array-based peptide libraries could be synthesized in a 384-well format. Also, MALDI-MS capabilities built into the platform to measure synthesis yields and purity for each array synthesis continue to improve to support more complex synthesis chemistry and library designs.

Putative epitope mapping was performed using the top peptide binding hits from the 3.3M full-coverage and 126K diverse libraries aligned to HER2. The epitope mapping results agreed with the known immunogens (Table 1). In addition, array binding assays performed in the presence and absence of HER2 identified those peptides most able to retain binding in the presence of the native antibody-HER2 interaction and highlighted conserved epitope positions. These assays also identified position-level substitution tolerance in the putative epitopes. Similar allowed epitope substitutions were observed in top binding hits from both the 3.3M full-coverage and 126K diverse libraries. Characterization of substitution tolerance at this position-level resolution could be important in antibody discovery to remove clones that lack specificity, have potential off-target effects and to select clones with pan-species or polymorphism reactivity. In addition to substitution tolerance, antibody binding tolerance with N-to-C and C-to-N sequence orientation was observed in at least one antibody clone which could be important in antibody discovery to identify clones with selectivity for one epitope orientation.

High-resolution epitope mapping and critical residue determination was performed using the top peptide binding hits identified from antibody binding in the presence and absence of HER2 using array-based 17K focused peptide libraries. These libraries had a sequence space focused on the putative epitopes identified from the 3.3M full-coverage library top binding peptides. For most clones, highly conserved epitope positions identified using the 17K focused library corroborate conserved motifs observed from the 3.3M full-coverage and 126K diverse libraries. Top binding peptides identified using the focused libraries contain more conserved epitope positions when compared to the 3.3M full-coverage and 126K diverse libraries. Many of the top peptides identified on the 17K focused library could compete at equivalence with the antibody-HER2 interaction in solution suggesting that these peptides represent much of the functional epitope.

In many cases, epitope mapping using array binding data generated with this platform can be done with widely available multiple sequence alignment algorithms like Clustal Omega or MUSCLE.^75, 76^ In this study, epitope mapping alignment performance was dependent on: 1) the level of top peptide motif conservation (specificity) of the antibody, 2) the library used to identify top binding peptides, and 3) the length of the immunogen region used for top peptide alignment. The 3.3M full-coverage library top peptide alignments could be performed using the full HER2 primary sequence for several of the antibodies measured, where the 126K diverse library top peptide alignments required restricting the alignment to the sequence region around the known immunogen. This immunogen alignment region restriction may or may not be a limitation in antibody discovery since the immunogen sequence is typically known, although epitope mapping accuracy with diverse library binding hits may decrease with increasing immunogen length. Also, custom alignment algorithms that can accommodate non-cognate flanking residues surrounding critical epitope residues in top peptides identified from a diverse peptide library may improve the performance of immunogen alignments for epitope mapping.

Taken together, these results highlighted key aspects of antibody binding on this platform and peptide library design for epitope mapping. High-resolution epitope and critical energetic residue identification on this platform showed that antibody clones selected with the same immunogen, even short peptide immunogens, can have very different functional epitopes. Also, top binding peptide results suggest that peptide library designs that maximize epitope information relative to total peptide length may improve high-resolution epitope mapping with peptide libraries. This could be explained by the fact that the entropic cost of an antibody binding a longer peptide must be overcome by an increase in favorable binding enthalpy across more peptide positions.

The epitope characterization methodology presented here is scalable approach to antibody epitope binning and mapping. On this platform, epitope mapping is performed first with potentially hundreds of clones. Second, clones are binned based on their identified epitopes without the requirement of a pairwise cross-blocking assay and the minimum N^2^/2 number of measurements for N clones. This allows epitope mapping to be completed much earlier in the discovery process with many more selected clones.

Broad and predictive antibody specificity characterization was also demonstrated using this peptide array platform. Top peptide target coverage metrics served as a measure of broad antibody binding specificity in a complex background matrix. Most notably, predictive capability of off-target interactions was demonstrated using the top antibody binding peptide hits from the 3.3M peptide full-coverage library and a proteome database search. Also important is the fact that antibody specificity is determined from the same array binding measurement as epitope mapping thereby multiplexing epitope and specificity measurements.

The antibody epitope and specificity characterization platform described here has the potential to improve antibody discovery productivity, reduce the risk of late-stage undesired side-effects, and identify rare clones with highly desired binding characteristics. This is achieved by providing information-rich, high-resolution, early-stage antibody characterization with the potential to scale to hundreds of clones. Ultimately the aim of this development is to produce an antibody characterization platform to identify panels of antibody leads that achieve high target epitope coverage, minimize off-target binding, and gain pan-species reactivity.

## Methods

### Library synthesis

Peptide libraries were chemically synthesized on 200 mm silicon oxide wafers functionalized with an aminosilane coating terminated with Boc-Glycine. For each cycle of the library synthesis, a layer of photoresist (containing a photoacid generator) was applied to the wafer, followed by exposure with UV light through the desired photomask – a general method which has been previously described.^71^ Light exposure activates the photoacid generator in the photoresist and allows for Boc-deprotection. Alignment of the photomask to the wafer prior to exposure allows for selected features to be deprotected, based on the openings of a given mask. Following UV light exposure, Boc removal was accelerated by heating, followed by removal of the photoresist. Application of the desired activated Boc-protected amino acid completes the coupling cycle. Repeating the cycles of deprotection and coupling with predetermined mask-amino acid combinations was used to achieve the desired peptide library. After completion of synthesis, each wafer was diced into 13 standard 75 × 25 mm microscope slides. Additionally, four MALDI-MS arrays were diced from the wafer. Slides were then subjected to a standard chemical cocktail to remove side chain protecting groups, and stored under dry nitrogen until use.

### MALDI mass spectrometry synthesis assessment

Two steps were performed to achieve high-resolution MALDI-MS of peptides *in situ* from the peptide arrays. First, the N-termini of peptides were labeled with a MALDI sensitivity enhancement reagent, Tris(2,4,6-trimethoxyphenyl)phosphine (TMPP). Second, the peptides were selectively and chemically cleaved from the surface without diffusion from the array features using gaseous ammonia.

For the matrix, Alpha-cyano-4-hydroxycinnamic acid (CHCA) was used along with two polymer standards, mono-dispersed PEG36 and PEG48-amine, which were doped into the matrix prior to application and were used as internal standards for calibration. The CHCA matrix was applied to the array surface using an automated sprayer (TM-Sprayer, HTX Technologies, LLC), providing a reproducible and uniform coating of matrix to the array surface.

MALDI mass spectra were acquired from array features using an Autoflex Speed MALDI MS (Bruker Daltonik), operating in positive ion reflectron mode, with ion source 1 at 19.00 kV, ion source 2 at 16.45 kV, reflector 1 at 20.95 kV, reflector 2 at 9.55 kV, lens at 7.9 kV, delay of 110 ns and signal suppression up to 400 Da. Prior to acquisition of the mass spectra, the target mass range (m/z = 400-4000) was internally calibrated using the two PEG-amine standards, with mass accuracy up to 0.01 Da.

Acquired mass spectra were imported into mMass V.5.5.0 software, smoothed (Gaussian method) and baseline subtracted. Desired peptides were identified based on the m/z of the peaks in the mass spectra, with a set tolerance of 0.5 Da and minimum signal-to-noise (S/N) of 3. Additional peaks, representing substitutions, deletions and/or by-products in the mass spectra were also identified if present (S/N >3). Desired peptide purity was calculated as a percent ratio from the peak intensities of the targeted peptide and the sum of the intensities of all identified peptides in the mass spectra. In addition, individual and average cycle efficiencies were calculated for all trimers, hexamers and other sequences as a percentage of the ratio between the peak intensities of the desired product peak and the sum of the desired plus undesired products, divided by the number of cycles. Verification that each array received the correct combinatorial synthesis was carried out by confirming the mass-to-charge ratio (m/z) of a set of protected hexamer peptides that represent each mask-amino acid combination at least once.

### Anti-HER2 monoclonal antibody panel

The panel of anti-HER2 monoclonal antibodies was sourced from multiple vendors. A single lot of each anti-HER2 monoclonal antibody from the full panel was used throughout the experiments on 17K, 125K, and 3.3M libraries. The following anti-HER2 clones were used within the anti-HER2 panel, grouped by vendor source: D8F12, 44E7, 29D8 (Cell Signaling 4290, 2248, 2165); A-2, C-3, C-12, SC-3B5, F-11 (Santa Cruz Biotechnology sc-393712, sc-377344, sc-374382, sc-33684, sc-7301); EP1045Y (Abcam ab1314182); 9G6 (LifeSpan Biosciences LS-150520); SPM495 (RayBiotech 119-13222); TF-3B5 (Thermo Fisher Scientific MA5-13675); RM228 (Origene TA352727); QO3B (Creative Diagnostics DCABH-2987); A24-V (MyBioSource MBS684037).

### HER2 competition array binding assay

Primary monoclonal antibodies were diluted in 120 μl of 1% mannitol (Sigma-Aldrich 63560) incubation buffer (10 g/l D-Mannitol, 1 PBS, 0.05% Tween 20, 0.1% Proclin 950, pH 7.4), to a final concentration of 1 nM and 0.25 nM then arranged in a 96-well Axygen plate (Corning). A 1:60 dilution of HER2 expressing whole cell lysate in RIPA buffer (Origene LY417979), or a lysate control (Origene LY500001), was spiked in as a competitor between replicate arrays. After a 5 minute pre-incubation of lysate and antibody at room temperature, 90 μl (17K or 126K libraries) or 600 μl (3.3M library) of primary antibody with competitor solution was transferred to array hybridization cassettes (Fig. S5) using a Bravo automated liquid handling platform (Agilent) with a 96-channel head. Cassettes were sealed and samples incubated for 30 minutes at 37 °C on a Teleshake 95 (Inheko). Arrays were then washed with PBST (1 PBS, 0.05% Tween 20, 0.1% Proclin 950, pH 7.4). Labeling of primary antibodies was performed as an additional incubation for 1 hour at 37 °C with a species-appropriate AF647-conjugated secondary antibody (Invitrogen A21235, A21245, A21445) in 1% mannitol incubation buffer. This was followed by a final wash series with PBST, then 18Ω ultrapure water. Arrays were then dried with 90% isopropyl alcohol (IPA).

Fluorescence intensity at each peptide was acquired using a 910 AL microarray scanner (Innopsys). Arrays were scanned with a 635 nm laser at 1 μm resolution for 17K and 126K arrays, or simultaneously scanned with 532 nm and 635 nm lasers at 0.5 μm resolution for 3.3M arrays.

### Sequence alignment and scoring for epitope binning, mapping and specificity

Multiple sequence alignments using the top 5 unique sequence hits were generated using ClustalW as provided by the R package msa.^77^ Alignments that display the target sequence included either the full target, or the listed immunogen sequence (Table 1), in the final alignment. Manual corrections to obvious ClustalW mis-alignment cases were done. A BLOSUM similarity scoring matrix (Fig. S2) was used to score alignments. The positional conservation score (*S_x_*) of an alignment used for putative epitope binning was defined as:

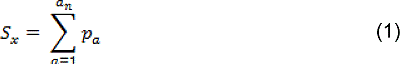

Where *a_n_* is the length of the amino acid alphabet used during library synthesis, *a* is a single amino acid within the set described by *a_n_*, and *p_a_* is the relative proportion of *a* at position *x* in the aligned set of peptide hits.

The positional coverage score (*C_x_*), and global coverage score (*C_glob_*) used as a specificity metric were defined as:

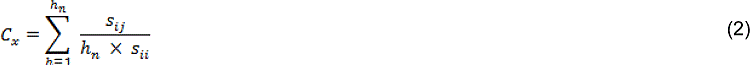

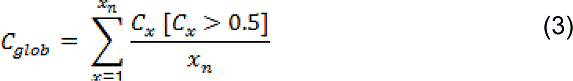

Where *h* is a peptide within a set of aligned hits of length *h_n_*, *s_ij_* is the BLOSUM similarity score for an amino acid at position with reference to the target HER2 immunogen sequence, *s_ii_* is the corresponding BLOSUM similarity score of the exact match to the target sequence, and *x_n_* is the total number of positions displayed within the alignment. The R package Logolas was used to display positional conservation and coverage scores as sequence logos above the original alignments for all clones across 17K, 126K, and 3.3M libraries.^78^

### Western Blot broad specificity assessment

Anti-HER2 monoclonal antibodies were used as primary antibodies in a series of Western Blots on cellular lysates of wild-type HEK293T (non-expressing HER2) cells (Origene LY500001) and HEK293T overexpressing HER2 (Origene LY417979). Lysates were directly loaded into each lane and Beta-Actin was used as a normalization control. Anti-HER2 antibodies were incubated with transfer blot membrane and labeled with species appropriate HRP-conjugated secondary antibody (Thermo 62-6520, Novex, A18805) and chromogenic HRP substrate (Fisher WP20004).

### Dot blot confirmation of array-predicted off-target binding partners

A nitrocellulose membrane was blocked with 5% BSA (Sigma-Aldrich A7906) in 1 x TBST (Fisher AAJ77500K2) for 1 hour after the direct addition of 1.5 μl of either recombinant HER4 (Sino Biological 10363-H08H), or recombinant ATF-6 Beta (Abnova H00001388-Q01). Wild-type HEK293T lysate (Origene LY500001) or HEK293T overexpressing HER2 (Origene LY417979) lysates were used as controls. Each antibody was tested at 0.4 μg/ml and 0.04 μg/ml in blocking buffer. Membranes were washed 3 times with TBST and incubated for 1 hour with a 1:2000 dilution of goat anti-mouse HRP (Thermo 62-6520) in blocking buffer for detection. Membranes were washed 3 times with TBST and visualized with chromogenic substrate. All steps were performed at room temperature. Quantitation of spot intensities was performed using ImageJ V2.0.0 software.

## Acknowledgements

We would like to acknowledge Tommy Armsby for his help with photolithographic mask and array assay cartridge design. We would also like to acknowledge Terrence Oneil for his supervision of all array syntheses. In addition, we would like to acknowledge Kathryn Nielsen and David Lomeli for their help with array binding assays and array imaging. Finally, we would like to thank Kathryn Sykes and Don Morris for their manuscript draft review.

