## Supplementary material for "A unified peptide array platform for antibody epitope binning, mapping, specificity and predictive off-target binding"

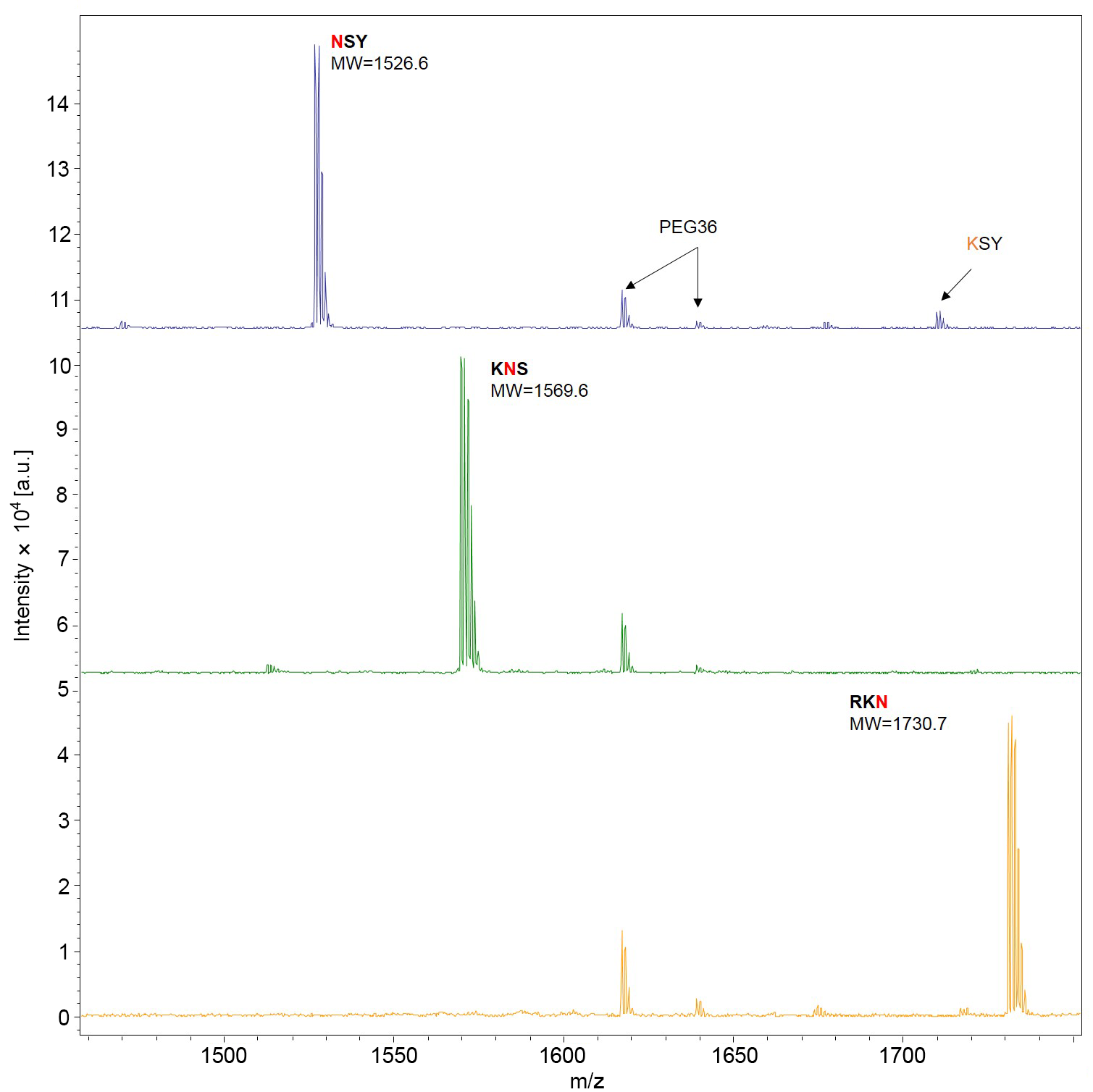

**Figure S1:** ***In Situ* MALDI-MS Overlapping Trimer Series from 3.3M Peptide Library Synthesis.** *In situ* MALDI mass spectra from three peptides in an overlapping trimer series, where asparagine (N) is the N-terminal (top), center (middle) and C-terminal (bottom) residue. This overlapping trimer MALDI-MS strategy is incorporated for all array synthesis steps providing quality assessment of every chemical cycle and nearest neighbor coupling effects. Full-length peptides (labeled with sequence and MW) contain three amino acids plus a tris (2,4,6-trimethoxyphenyl)phosphonium-acetyl (TMPP-Ac) group and a 6-unit polyethylene glycol (PEG) linker. A single amino acid substitution side-product (KSY, top mass spectra) is marked by an arrow. Mono-dispersed PEG-amine (PEG36) was added in the MALDI matrix as a standard for calibration. Signals from [M+H] and [M+Na] PEG36-amine are present in each mass spectra at m/z=1616.9 and 1639.0 respectively.

**Table S1:** Overlapping hexamer peptide series *In Situ* MALDI-MS Photolithographic Synthesis Verification for 3.3MPeptide Array Library

| ID | Expected Product Sequence | Mass-to-Charge (m/z) | Product Peak Intensity | Signal-to-Noise (S/N) | Synthesis Verification |
| --- | --- | --- | --- | --- | --- |
|  |  |  |  |  | S/N>10 |
| 175 | YWPLVG | 1787.84 | 27,884 | 114.7 | PASS |
| 176 | KNSYWP | 2105.84 | 31,234 | 104.3 | PASS |
| 177 | DHRKNS | 2278.87 | 21,270 | 103.9 | PASS |
| 178 | LVGDHR | 1960.87 | 18,461 | 103.3 | PASS |
| 179 | YWPLVG | 1787.84 | 35,007 | 132.5 | PASS |
| 180 | KNSYWP | 2105.84 | 38,735 | 131.5 | PASS |
| 181 | DHRKNS | 2278.87 | 20,476 | 98.3 | PASS |
| 182 | LVGDHR | 1960.87 | 12,707 | 71.3 | PASS |
| 183 | YWPLVG | 1787.84 | 40,130 | 131.4 | PASS |
| 184 | KNSYWP | 2105.84 | 40,429 | 115.5 | PASS |
| 185 | DHRKNS | 2278.87 | 17,863 | 92.3 | PASS |
| 186 | LVGDHR | 1960.87 | 21,731 | 108.5 | PASS |
| 187 | YWPLVG | 1787.84 | 37,461 | 142.8 | PASS |
| 188 | KNSYWP | 2105.84 | 34,774 | 115.0 | PASS |
| 189 | DHRKNS | 2278.87 | 28,735 | 132.7 | PASS |
| 190 | LVGDHR | 1960.87 | 24,062 | 134.6 | PASS |
| 191 | YWPLVG | 1787.84 | 47,601 | 184.6 | PASS |
| 192 | KNSYWP | 2105.84 | 39,256 | 116.9 | PASS |
| 193 | DHRKNS | 2278.87 | 14,122 | 86.7 | PASS |
| 194 | LVGDHR | 1960.87 | 16,244 | 88.9 | PASS |
| 195 | YWPLVG | 1787.84 | 57,577 | 193.0 | PASS |
| 196 | KNSYWP | 2105.84 | 44,658 | 152.5 | PASS |

|  | **A** | **R** | **N** | **D** | **C** | **Q** | **E** | **G** | **H** | **I** | **L** | **K** | **M** | **F** | **P** | **S** | **T** | **W** | **Y** | **V** |  |
| --- | --- | --- | --- | --- | --- | --- | --- | --- | --- | --- | --- | --- | --- | --- | --- | --- | --- | --- | --- | --- | --- |
| **A** | 6 |  |  |  |  |  |  |  |  |  |  |  |  |  |  |  |  |  |  |  | **A** |
| **R** | -1 | 5 |  |  |  |  |  |  |  |  |  |  |  |  |  |  |  |  |  |  | **R** |
| **N** | -2 | 0 | 6 |  |  |  |  |  |  |  |  |  |  |  |  |  |  |  |  |  | **N** |
| **D** | -2 | -2 | 6 | 6 |  |  |  |  |  |  |  |  |  |  |  |  |  |  |  |  | **D** |
| **C** | 0 | -3 | -3 | -3 | 9 |  |  |  |  |  |  |  |  |  |  |  |  |  |  |  | **C** |
| **Q** | -1 | 1 | 4 | 0 | -3 | 5 |  |  |  |  |  |  |  |  |  |  |  |  |  |  | **Q** |
| **E** | -1 | 0 | 6 | 4 | -4 | 2 | 5 |  |  |  |  |  |  |  |  |  |  |  |  |  | **E** |
| **G** | 6 | -2 | 0 | -1 | -3 | -2 | -2 | 6 |  |  |  |  |  |  |  |  |  |  |  |  | **G** |
| **H** | -2 | 0 | 1 | 6 | -3 | 0 | 0 | -2 | 8 |  |  |  |  |  |  |  |  |  |  |  | **H** |
| **I** | -1 | -3 | -3 | -3 | -1 | -3 | -3 | -4 | -3 | 4 |  |  |  |  |  |  |  |  |  |  | **I** |
| **L** | -1 | -2 | -3 | -4 | -1 | -2 | -3 | -4 | -3 | 4 | 4 |  |  |  |  |  |  |  |  |  | **L** |
| **K** | -1 | 2 | 0 | -1 | -3 | 1 | 1 | -2 | -1 | -3 | -2 | 5 |  |  |  |  |  |  |  |  | **K** |
| **M** | -1 | -1 | -2 | -3 | -1 | 0 | -2 | -3 | -2 | 5 | 5 | -1 | 5 |  |  |  |  |  |  |  | **M** |
| **F** | -2 | -3 | -3 | -3 | -2 | -3 | -3 | -3 | -1 | 0 | 0 | -3 | 0 | 6 |  |  |  |  |  |  | **F** |
| **P** | -1 | -2 | -2 | -1 | -3 | -1 | -1 | -2 | -2 | -3 | -3 | -1 | -2 | -4 | 7 |  |  |  |  |  | **P** |
| **S** | 1 | -1 | 1 | 0 | -1 | 0 | 0 | 0 | -1 | -2 | -2 | 0 | -1 | -2 | -1 | 4 |  |  |  |  | **S** |
| **T** | 0 | -1 | 0 | -1 | -1 | -1 | -1 | -2 | -2 | -1 | -1 | -1 | -1 | -2 | -1 | 5 | 5 |  |  |  | **T** |
| **W** | -3 | -3 | -4 | -4 | -2 | -2 | -3 | -2 | -2 | -3 | -2 | -3 | -1 | 4 | -4 | -3 | -2 | 11 |  |  | **W** |
| **Y** | -2 | -2 | -2 | -3 | -2 | -1 | -2 | -3 | 2 | -1 | -1 | -2 | -1 | 6 | -3 | -2 | -2 | 4 | 7 |  | **Y** |
| **V** | 0 | -3 | -3 | -3 | -1 | -2 | -2 | -3 | -3 | 4 | 4 | -2 | 5 | -1 | -2 | -2 | 0 | -3 | -1 | 4 | **V** |
|  | **A** | **R** | **N** | **D** | **C** | **Q** | **E** | **G** | **H** | **I** | **L** | **K** | **M** | **F** | **P** | **S** | **T** | **W** | **Y** | **V** |  |

**Figure S2: BLOSUM Consensus and Alignment Scoring Matrix.** The BLOSUM matrix used for the calculation of top peptide array hit consensus and epitope coverage scores across all anti-HER2 clones on 17K, 126K, and 3.3M libraries.

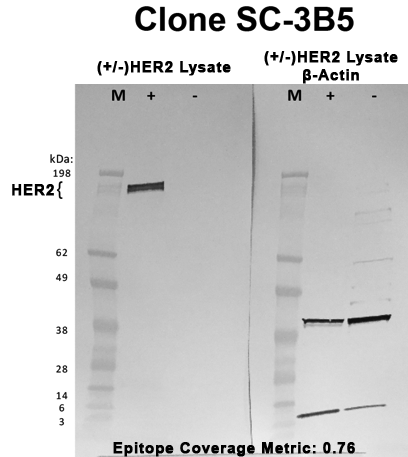

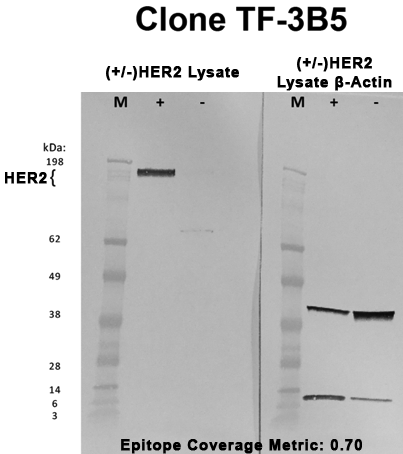

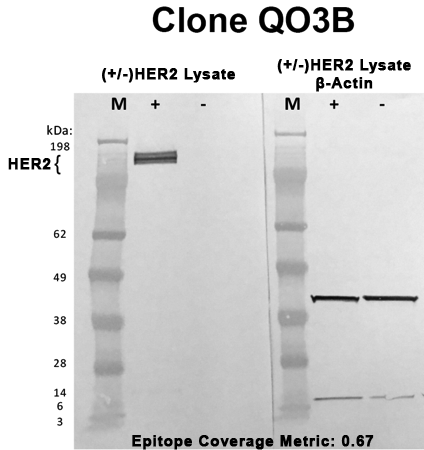

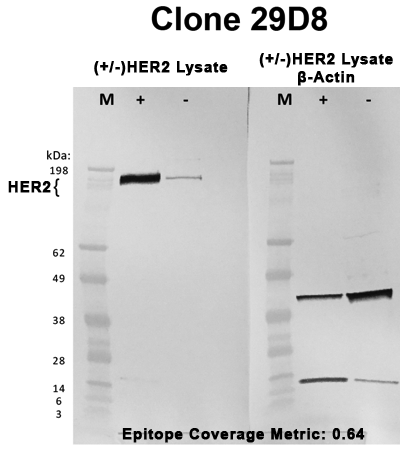

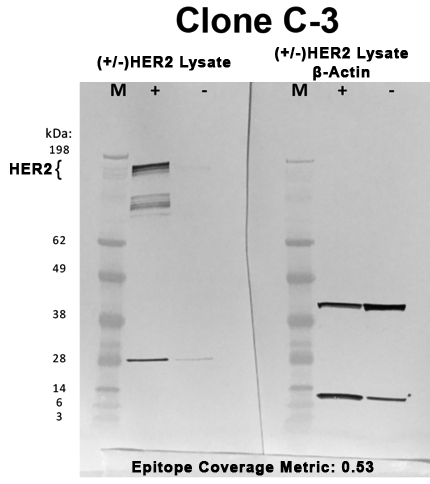

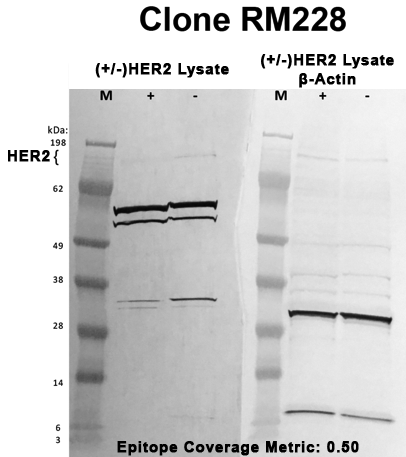

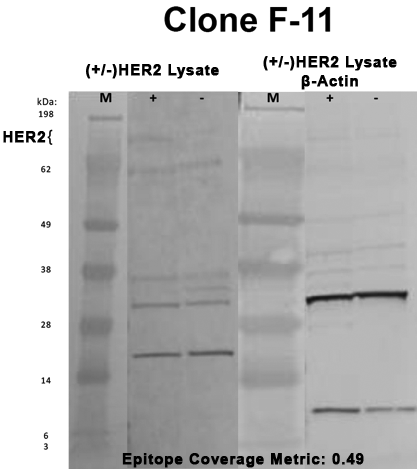

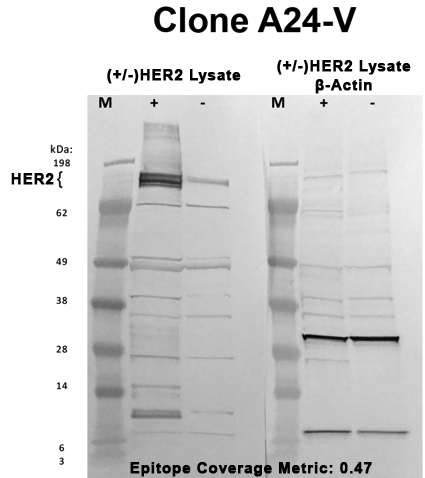

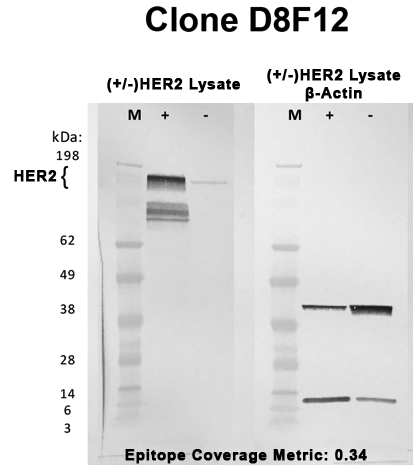

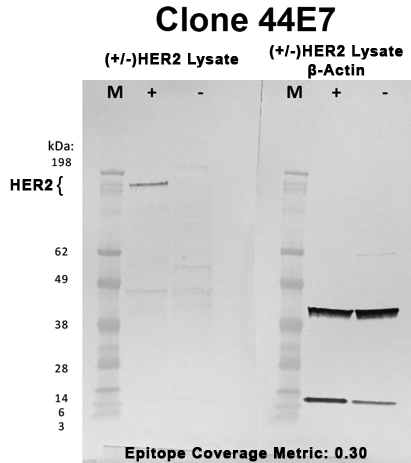

**Figure S3: (+/-)HER2 Expression HEK293T Cell Lysate Western Blot mAb Broad Binding Specificity Characterization.** Anti-HER2 clone Western Blots with (+/-)HER2 cell lysate on left half of gel images, β-actin control on right half of gel images. Images are sorted by decreasing epitope coverage metric (Table 3, Main Text)

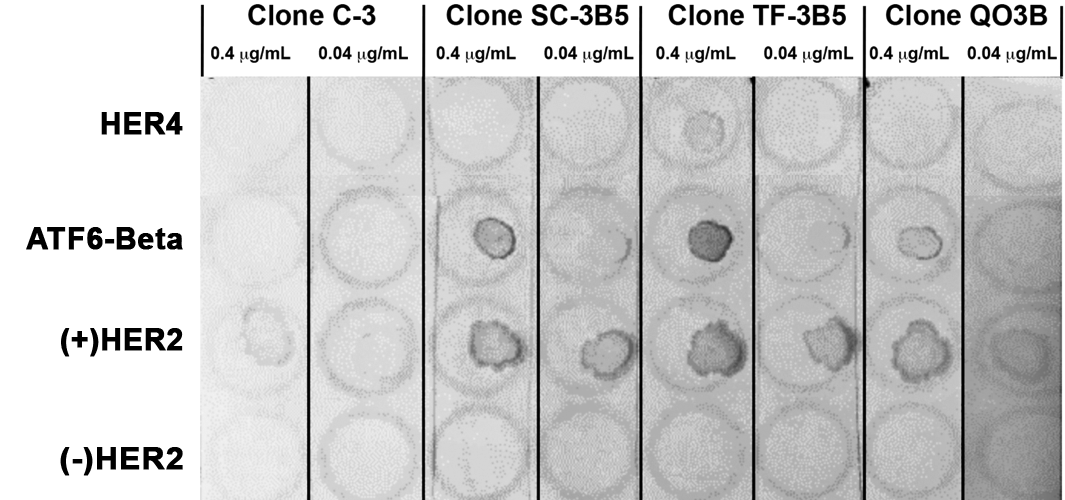

**Figure S4: Dot-blot Binding Assays for Array-Predicted Off-Target Interaction Partners.** Dot-blot binding results with clones C-3, SC-3B5, TF-3B5, and QO3B HER2 and 3.3M peptide array predicted off target binding partners. Antibody clones at two concentrations shown in columns and predicted off target purified recombinant proteins HER4, and ATF-6 Beta shown in rows. (+/-)HER2 overexpression HEK293T lysate controls shown in last two rows.

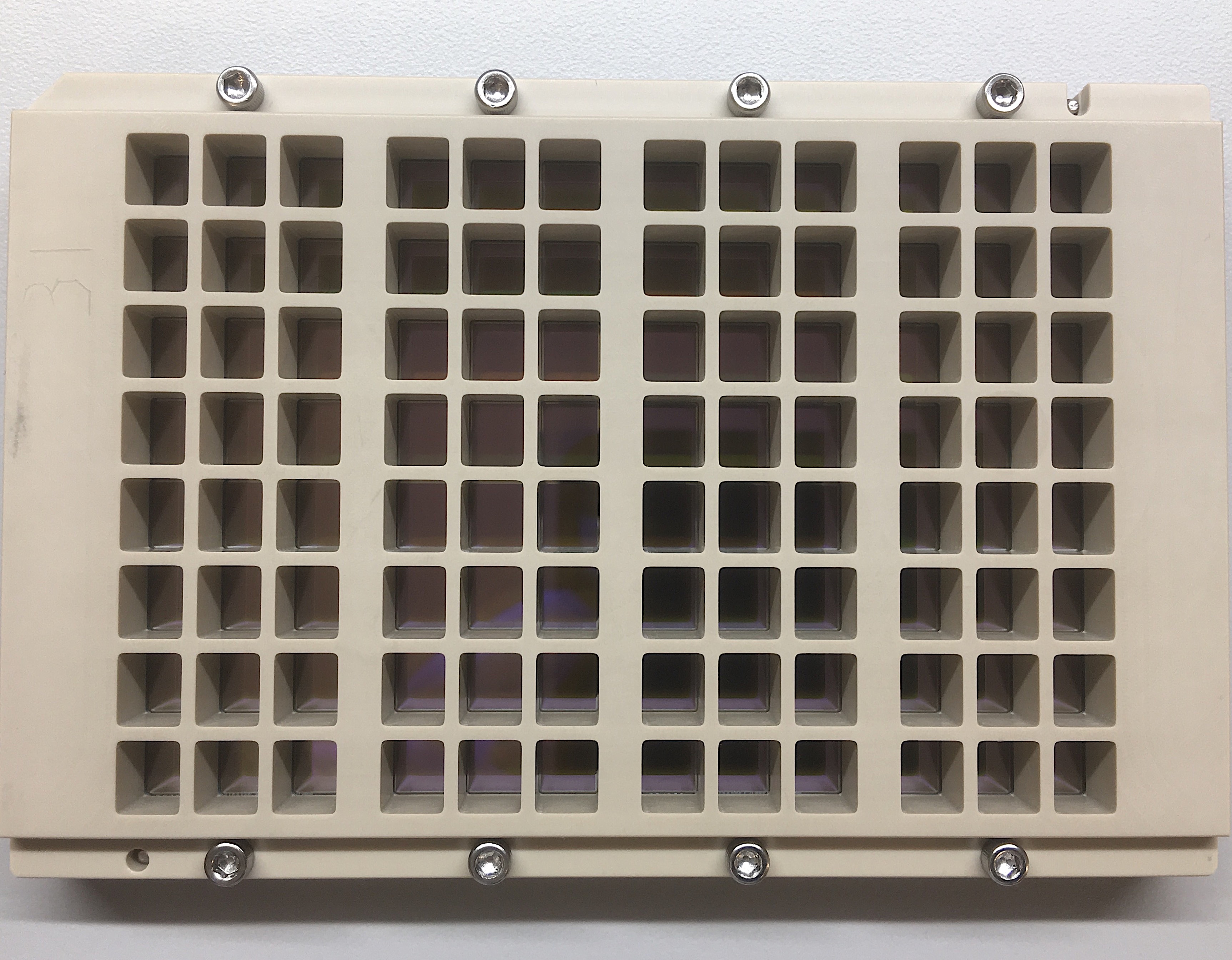

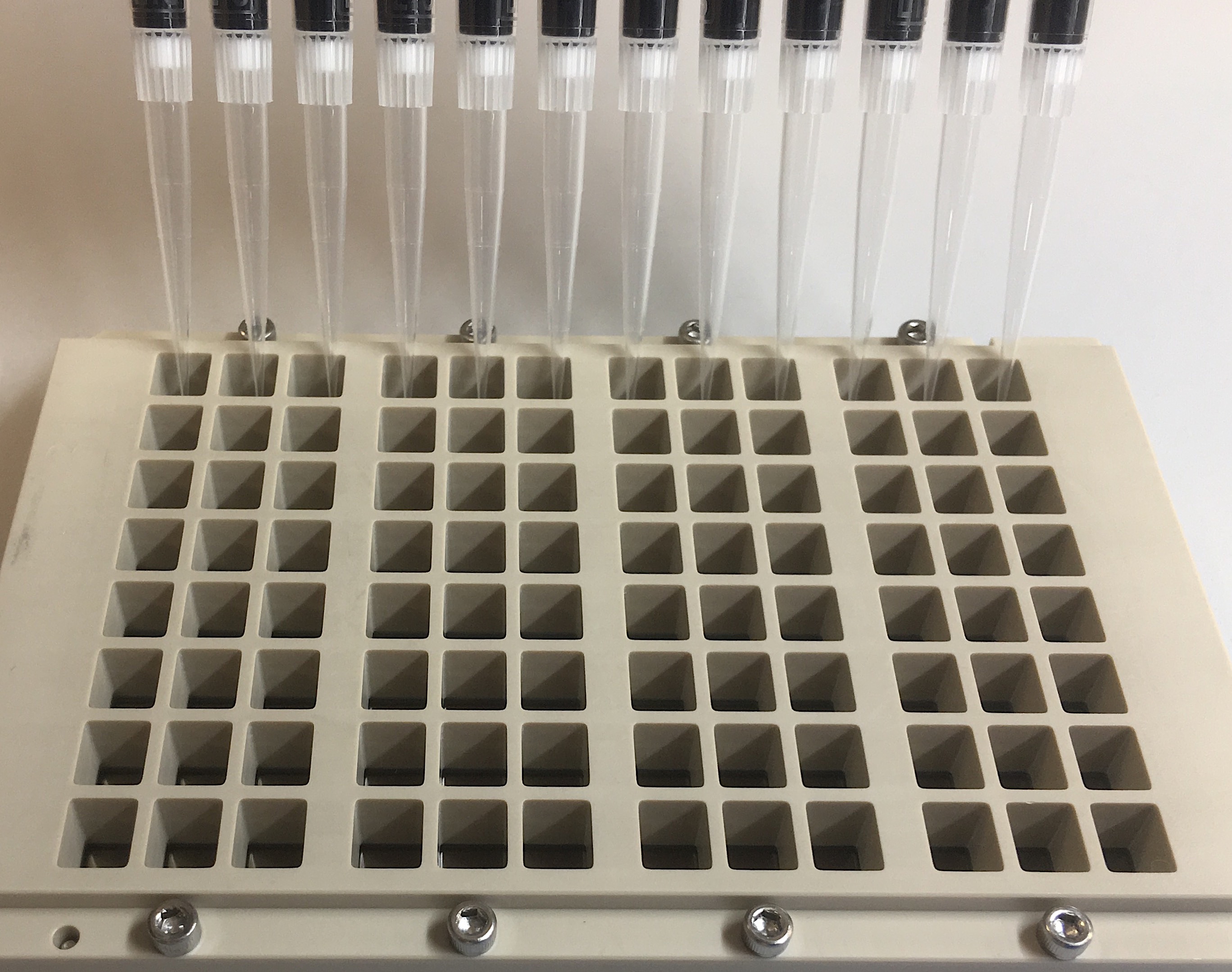

**Figure S5: 96-Well Binding Assay Cassette for 126K Diverse and 17K Focused Peptide Arrays.** Each well contains an entire peptide array. The cassette conforms to SLAS microplate standards and is compatible with common automation and liquid handling hardware.
